# Reactivation during sleep segregates the neural representations of episodic memories

**DOI:** 10.64898/2026.04.08.717230

**Authors:** Sarvia Aquino Argueta, Andrew Lazarus, Fiona Yao, Charli Taylor, Laura K Shanahan, Kenneth A Norman, Lila Davachi, Thorsten Kahnt, Ken A Paller, Eitan Schechtman

## Abstract

Sleep involves the reactivation of recently acquired memories, thereby shaping the neural representations supporting them. An essential feature of episodic memory (i.e., memory for events) is the link between specific elements (e.g., a cake) and the context in which they were embedded (e.g., a birthday party). We investigated how reactivation during sleep impacts item-context binding. Participants (*N* = 22) formed stories linking together objects in unique contexts. Using functional MRI, we measured the overlap between neural representations for contextually linked objects. During a 90-min nap, some object memories were reactivated by unobtrusively presenting object-specific sounds. Across multiple brain regions, reactivation reduced the overlap between representations of contextually linked memories, promoting object specificity. Furthermore, reactivation reduced the representational overlap between contexts, thereby promoting context specificity. Taken together, data suggest that reactivation during sleep decontextualizes memories. These results inform our understanding of how sleep contributes to interlinked memories embedded in naturalistic contexts and demonstrate that memory reactivation during sleep changes representations of memories within their distinctive contexts.

## Introduction

Episodic memories (memories for specific events) reflect information on the three Ws of the event: What happened, when it happened, and where it happened ^1,2^. This type of memory is contextual in nature; to tell one memory from another, such as where you parked your e-bike this morning vs yesterday, the specific context must be reinstated and distinguished from similar competing contexts^1^. Over time, episodic memories evolve and transform: some details are lost, but the semantic elements are preserved in the long term. The process of memory transformation through decontextualization and semantization is thought to manifest through forgetting, generalization, gist extraction, and schema creation^3^. On the neural level, semantization is thought to be supported by a gradual shift in memory representations: from strong hippocampal dependency to reliance primarily on the neocortex^4,5^.

The process through which memory representations at the systems level transform over time is termed systems consolidation. Research on consolidation has considered both behavioral manifestations (e.g., changes in retention) and physiological manifestations (e.g., changes in neural representations), over time spans ranging from minutes^6^ to years^7^. The Complementary Learning Systems model^5^ suggests that consolidation occurs through reactivation of hippocampal traces that gradually mold cortical circuits, and proposes that this process occurs during periods of sleep^8–11^. In support of sleep’s role in consolidation, a multitude of studies have shown that sleep improves memory retention for various tasks^12,13^. Furthermore, several studies have found evidence for sleep’s contribution to memory transformation (e.g., generalization and gist abstraction), supporting the predictions of the Complementary Learning Systems model^14–16^.

Critically, changes in representation, rather than retrieval probabilities, are central to theories of consolidation. Several studies have examined how consolidation impacts neural representation over several months in humans (e.g., ^17,18^). Whereas many of these studies considered memory trace transformation broadly, some have focused primarily on the concept of decontextualization, finding that consolidation leads to reorganization and semantization of memories over time ^19,20^. Despite the hypothesized importance of sleep in this process and the abundance of evidence on sleep’s behavioral effects, sleep’s role in representational transformation of episodic memories has received less attention. In a recent study, Cowan and colleagues have demonstrated that representations of interrelated memories in the hippocampus change over a period of sleep: representations in the anterior hippocampus become less overlapping, whereas the opposite pattern appears in the posterior hippocampus^21^. Furthermore, the density of sleep spindles, a sleep-specific electrophysiological waveform that has been linked to consolidation^22^, was positively correlated with both changes in hippocampal-cortical functional connectivity and representational overlap in the ventromedial prefrontal cortex^23^.

Although these findings support the key role of sleep in memory transformation, they fail to directly address the central role of sleep-based reactivation in memory consolidation. Furthermore, conclusions are limited because of their dependence on correlational findings across subjects, leaving open several competing interpretations such as reverse causality (i.e., changes in representation impacting sleep rather than the other way around) and broader interindividual differences that extend beyond spindle density (which itself correlates with traits such as learning ability and other metrics of intelligence^24^). Furthermore, the changes observed do not directly implicate the process of reactivation during sleep, which has been hypothesized to be key to consolidation. In this study, we used targeted memory reactivation (TMR), a non-invasive technique developed to causally bias consolidation during sleep by unobtrusively presenting learning-related stimuli^25,26^. Most TMR studies have demonstrated its effects on memory retention^27^, whereas fewer have examined its effects on neural representations. Berkers and colleagues^28^ showed that TMR using sound cues increased the integration of the occipital cortex with memory-related networks, although this effect was not specific to cued memories. More recently, Rakowska and colleagues^29^ found that TMR increased activity in the left precuneus selectively for cued memories, and this activation was linked with subsequent performance in a sensorimotor task. Importantly, neither those studies nor others examining changes in neural activity (e.g., ^30–32^) have directly examined changes in neural representation, whether related to decontextualization or otherwise.

To explore this question, we designed a within-subject study involving EEG, functional MRI (fMRI), and TMR (Figure 1). Participants were first instructed to form stories incorporating four objects (e.g., animals, household items, musical instruments), thus creating multiple rich episodic contexts that include many of the elements that characterize real-life contexts (e.g., a causal relationship between elements, a shared space, a shared theme, overlapping temporal dynamics). These contexts therefore consisted of both a shared thematic/semantic core and episode-specific elements such as a shared imagined time and place. The neural representations of each object were extracted using multivariate pattern analysis applied to fMRI data. Similarity between representations of objects within the same story served as a proxy for item–context binding. Over a 90-minute nap, participants were unobtrusively exposed to sounds congruently related to some of the objects. The representations were then examined again, and changes in the contextual overlap were assessed for both cued and non-cued contexts, with decreases in similarity values reflecting decontextualization. Furthermore, we considered how reactivation during sleep changed between-context differentiation by examining changes in overlap between objects associated with different contexts. Our results suggest that reactivation during sleep leads to decontextualization of memory traces and increases dissimilarity between distinct contexts.

**Figure 1.**
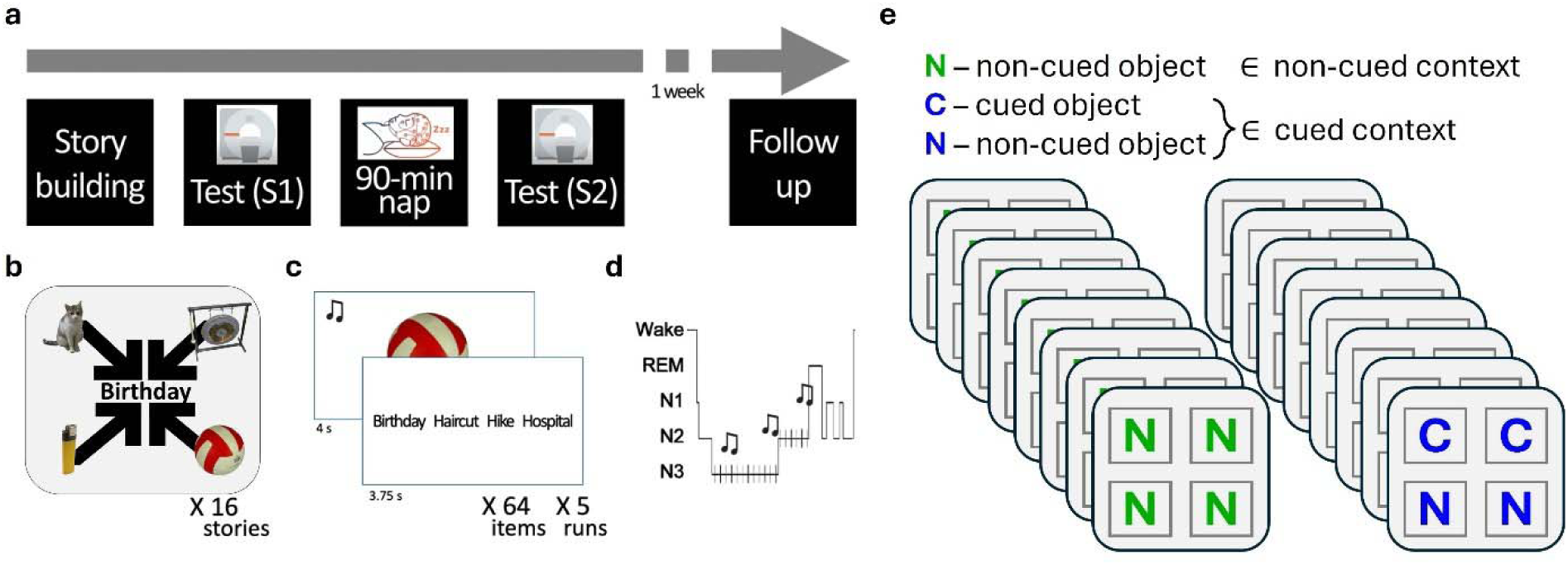
Experimental Design. (a) Outline of procedure. Images indicate whether EEG or MRI data were recorded. (b) Participants (N = 22) were required to build 16 stories, each centered on a specific scenario (e.g., attending a birthday party) and incorporating four objects in their stories. Stories were then overtrained to reach ceiling memory levels. (c) After memorizing the stories, they were tested inside the MRI scanner (i.e., Retrieval Task). Across five runs, each image was presented along with its congruent sound, and participants were instructed to recall which story it went with. (d) Outside of the scanner, participants were fitted with an EEG cap and given the opportunity to nap for 90 minutes. During the nap, sound cues linked with objects were unobtrusively presented during Non-REM sleep. After the nap, participants completed the Retrieval Task again in the scanner. (e) Of the 16 stories constructed (i.e., contexts), objects from eight stories were reactivated during sleep. Altogether, sounds for half the objects in half the stories were presented during sleep. Images taken from the Bank of Standardized Stimuli33.

## Results

Participants formed 16 stories, with each story linking four different objects together in a cohesive narrative (Figure 1b). Then, in the MRI scanner, participants were presented with object images and their congruent sound and had to retrieve the associated context (i.e., story; Figure 1c). During a subsequent afternoon nap, sounds associated with some of the objects were unobtrusively presented, putatively reactivating these memories and leading to memory consolidation (Figure 1d). In half of the contexts, half of the objects were cued (Cued ∈ Cued Context), and the other were not (Non-cued ∈ Cued Context). In the other half of contexts, no objects were cued (Non-Cued ∈ Non-Cued Context; Figure 1e). After the nap, participants returned to the scanner and completed the same task as before.

### Memory performance was at ceiling across multiple tests

Since the study focused on the transformation of neural representations rather than changes in memory retrievability, participants were overtrained on item-context associations during initial learning. To quantify recall accuracy, we calculated the percentage of correctly recalled item-context associations during the Retrieval Task for each condition across both sessions (Table 1). A repeated-measures 2 X 2 ANOVA (session, cued vs non-cued contexts) revealed a main effect of session, indicating improved accuracy across time (*F*(1,21) = 10.09, *p* = 0.005). It is unclear whether this improvement was due to practice, a testing effect^34^, the effect of sleep, or some combination of these factors. There were no significant main effects of cuing and no interaction effect (*F*(1,21) = 1.49, *p* = 0.24; *F*(1,21) = 1.53, *p* = 0.23). Similar effects were obtained when considering the cued and non-cued objects separately (2 x 3 repeated measures ANOVA; main effect of session, *F*(1,21) = 12.4, *p* = 0.002; no main effect of cuing, *F*(2,42) = 0.92, *p* = 0.41; no interaction, *F*(2,42) = 0.81, *p* = 0.45). These findings suggest that while memory performance improved across sessions, cuing did not produce a measurable behavioral effect, likely due to near-ceiling performance across all conditions.

**Table 1.** Behavior Results: Performance in test % correct (± SEM).

|  | Cued Context | Non-Cued Context |
| --- | --- | --- |
| **Pre-Sleep (Session 1)** | 93.07% ± 1.43% | 94.37% ± 1.59% |
| **Post-Sleep (Session 2)** | 97.66% ± 0.54% | 97.73% ± 0.53% |

Memory tests run after the scan confirmed that memory was at ceiling, and remained so for at least one week. Immediately after Session 2, participants were instructed to list all objects per context in the Recall Task and performed at high levels (97.44% ± 0.65%; range: 90.62%-100%). Approximately one week later, participants completed a Recognition Task, demonstrating that their memory for the objects remained high (hit rate: 95.24% ± 0.81%; range: 85.94%-100%; false alarms: 1.63% ± 0.48%; range: 0%-7.81%). Similarly, performance was high in the Sorting Task, where participants had to link each context with its four objects (97.09% ± 1.55%; range: 65.63%-100%).

### Reactivation during sleep reduced object-context binding

The study aimed to examine the effects of reactivation during sleep on the neural representations of objects within their context. To examine this, we identified the multi-voxel representations of the 64 objects and examined similarities among contextually bound objects both before and after sleep (Figure 2a). This approach was first applied across all regions of the posterior medial network (considered jointly as a single region of interest), a subnetwork of the default mode network that has been shown to represent contexts^35,36^ (Figure 2b, left). We first used data from Session 1 to test whether these regions represent context in our task. To do this, we compared the correlations among contextually bound objects with the correlations among objects from different contexts. Results show that the representations among contextually bound objects were more correlated than the representations of objects linked with different contexts, demonstrating that the posterior medial network is sensitive to contexts in our task (*t*(21) = 3.27, *p* = 0.004). Next, we examined the effects of sleep on correlations within contexts. GLM-derived β values for each object were correlated across voxels with those of the three objects of the same context. Correlation values were z-transformed and averaged across context for both cued and non-cued contexts. A 2 x 2 repeated measures ANOVA (session, cued vs non-cued contexts) found no main effects of session (*F*(1,21) = 0.01, *p* = 0.91) or cuing (*F*(1,21) = 0.56, *p* = 0.46). Critically, however, there was a significant interaction (*F*(1,21) = 5.05, *p* = 0.035). This interaction was driven by higher post-sleep correlation values for non-cued contexts relative to cued contexts (*t*(21) = 2.21, *p* = 0.039; Figure 2b). Similarly, comparing changes in correlation values across sessions revealed increased correlations for the non-cued contexts relative to the cued ones (*t*(21) = 2.25, *p* = 0.035; Figure 2c). Supplementary Figure 1 shows data for brain regions linked with the posterior medial network, as well as other related brain regions.

**Figure 2.**
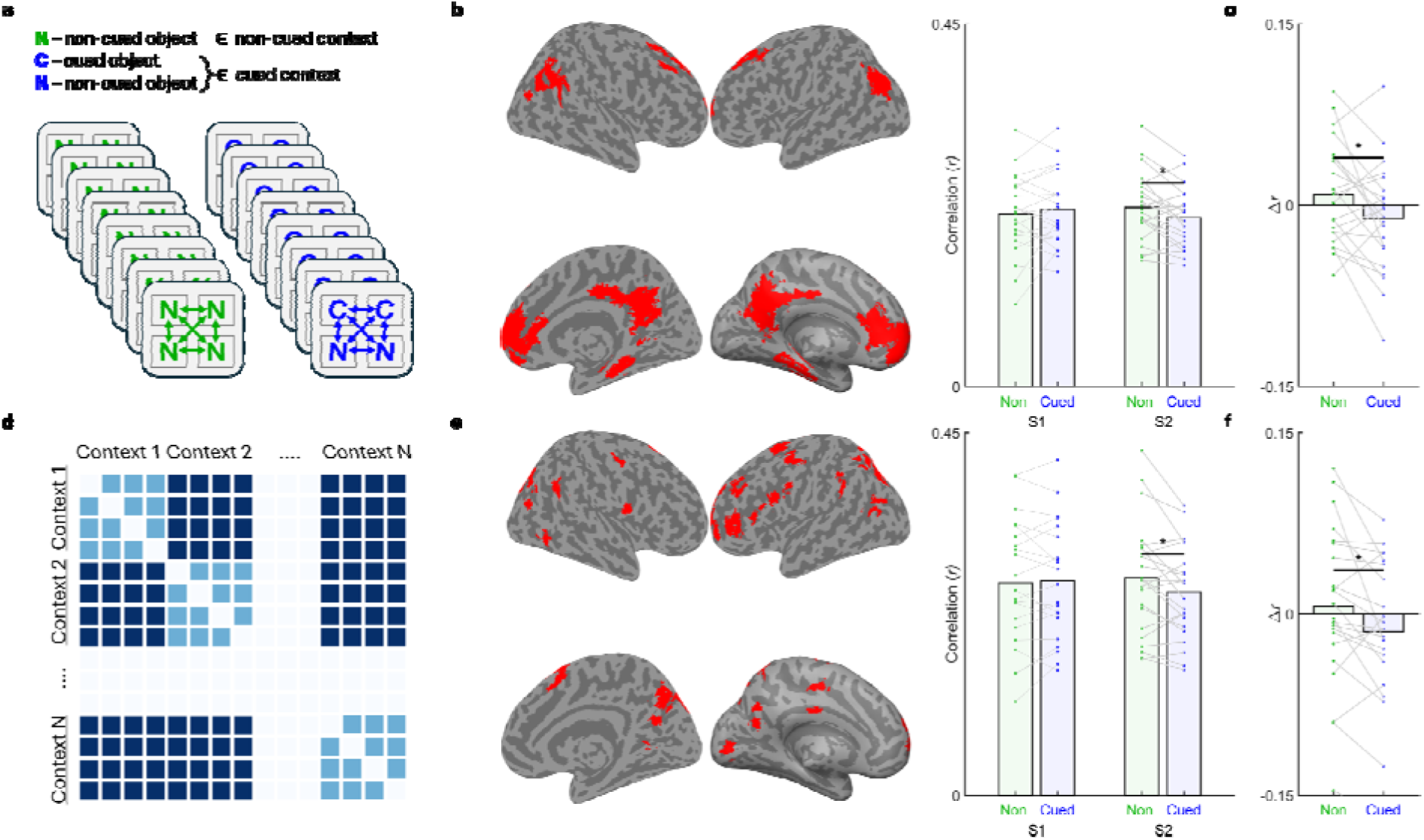
– Reactivation during sleep reduced object-context binding. (a) Analyses compared correlations among contextually bound objects within contexts that were cued (i.e., contexts which included objects which were reactivated during sleep) or non-cued (objects marked in blue and green, respectively). (b) Examining the posterior medial network in its entirety (regions of interest marked in red), reactivation reduced within-context correlations in cued contexts relative to non-cued contexts. (c) Differences over sleep between cued and non-cued contexts. (d) To complement analyses on the posterior medial network, we used a data-driven approach to identify cortical regions that reflect object-context associations. This analysis identified regions in which within-context correlations (light blue squares) were higher than across-context correlations (dark blue squares) in Session 1. (e) Examining the identified regions (marked in red) together, they show similar patterns of results, with reduced within-context correlations following reactivation. (f) Differences over sleep between cued and non-cued contexts. S1 – Session 1 (pre-sleep); S2 – Session 2 (post-sleep). Lines represent individual participants. * p < 0.05.

To examine the inner dynamics within cued contexts, we calculated the cross-session change in correlations between the two cued objects and between the two non-cued objects within the cued context (Blue Cs and Blue Ns in Figure 2a, respectively). Cued objects showed a trend toward decreased correlations when compared with objects in the non-cued contexts (*t*(21) = 1.83, *p* = 0.08), whereas the non-cued objects within the cued contexts showed significantly decreased correlations when compared with objects in the non-cued contexts (*t*(21) = 2.54, *p* = 0.019). Changes in correlation values within cued and within non-cued objects in cued contexts were not significantly different (*t*(21) = 0.69, *p* = 0.5). Taken together, these results indicate that cuing reduced the overlap in the neural representation of contextually bound objects, supporting the hypothesis that reactivation during sleep serves to decontextualize memories by weakening object-context links.

Although the posterior medial network has been shown to represent contexts in which memories reside^35,36^, we complemented our region-of-interest-driven analyses with data-driven analyses using a searchlight approach. The goal of these analyses was to confirm that the effects observed in the posterior medial network are similarly observed when considering areas that are sensitive to contextual binding in our task. Using data collected in Session 1, we identified which of 1,000 cortical parcels^37^ represent contextual binding by statistically comparing within-to between-context correlations (*p* < 0.05, FDR-corrected; Figure 2d). A complementary analysis using a whole-brain searchlight did not reveal any subcortical regions that crossed statistical thresholds (Supplementary Figure 2). Both data-driven analyses revealed multiple cortical regions that reflected contextual binding between objects (i.e., showed higher within-vs between-context correlations), including the superior and inferior parietal lobule, the lateral prefrontal cortex, and the precuneus cortex (Figure 2e; Supplementary Figure 2).

All identified parcels were submitted to the same analysis described above (as a single region of interest). A 2 x 2 repeated measures ANOVA (session, cued vs non-cued contexts) found no main effects of session (*F*(1,21) = 0.1, *p* = 0.76) or cuing (*F*(1,21) = 2.31, *p* = 0.14), mirroring the results obtained for the posterior medial network. As before, a significant interaction between session and cuing was observed (*F*(1,21) = 7.2, *p* = 0.014). This was driven by lower post-sleep correlations within the cued contexts relative to the non-cued contexts (*t*(21) = 2.47, *p* = 0.022; Figure 2e). Similarly, examining the change in correlation across sessions showed increased correlations for the non-cued contexts relative to the cued ones (*t*(21) = 2.68, *p* = 0.014; Figure 2f). When examining the cued and non-cued objects within cued contexts, the result for the posterior medial network were replicated: cued objects showed a trend toward decreased correlations relative to the non-cued contexts (*t*(21) = 1.85, *p* = 0.08), non-cued objects showed significantly decreased correlations relative to the non-cued contexts (*t*(21) = 2.51, *p* = 0.02), and the two types of objects did not differ from each other (*t*(21) = 0.5, *p* = 0.63). The results for the two networks converge and suggest that across brain regions representing contexts, reactivation during sleep segregates memories’ neural representations from one another.

### Reactivation leads to transformation in the neural representation of individual objects

As a complementary analysis, we examined whether reactivation led to a transformation of the neural representation of individual objects between sessions, as shown in a previous study using EEG^38^. Throughout this study, we operationalized the term “object representation” using a broad definition, i.e., any neural activity that is object-specific, as derived from the GLM. This includes, by definition, the neural correlates of item-context binds for a specific object. A transformation of an object’s representation and a change in item-context binds are therefore partially overlapping processes. A change in an object’s neural representation can lead to it being less similar to its contextual counterparts, but it could also lead to it being equally similar or more similar to them. To disentangle this complex relationship, we examined the cross-session correlations between object representations. First, we asked whether reactivation during sleep changes the correlation between the session-1 representation of an object and its session-2 representation. Indeed, we found that objects in cued contexts were less correlated to themselves relative to objects in non-cued contexts (posterior medial network: *t*(21) = 2.09, *p* = 0.049; regions identified using the data-driven approach: *t*(21) = 2.17, *p* = 0.041). Lower cross-session correlations indicate that reactivation transformed objects, rendering them less similar to their pre-sleep baseline.

These results suggest that, indeed, reactivation led to representational changes, but they leave open the question of whether individual-object changes are the main driver of the observed changes in the neural correlates of object-context binds. To test this, we calculated the cross-session similarities not only for each object with itself, but also with its contextual counterparts (e.g., if objects A and B were contextually linked, the analysis would include the correlation between object A’s representation in session 1 and object B’s representation in session 2 and vice versa). Correlations were submitted to a 2 X 2 ANOVA (correlation with same object vs correlation with other contextually bound objects; cued context vs non-cued context). In this analysis, an interaction driven by a higher-magnitude change in the representation fidelity of an object would suggest that objects are transformed above and beyond the transformation observed for item-context binds. Results revealed a significant main effect of same-object vs. different-object, demonstrating that object representations were better correlated across sessions within themselves than with other objects, as is to be expected (posterior medial network: *F*(1,21) = 7.91, *p* = 0.01; regions identified using the data-driven approach: *F*(1,21) = 11.52, *p* = 0.003). Additionally, results revealed a main effect of condition, with lower cross-session correlations for objects in cued contexts relative to objects in non-cued contexts (posterior medial network: *F*(1,21) = 7.45, *p* = 0.01; regions identified using the data-driven approach: *F*(1,21) = 4.39, *p* = 0.048). Although these results aren’t completely independent of the results reported in previous paragraphs (calculated for each session separately), they converge to demonstrate that reactivation segregates objects from their contextually bound counterparts. Critically, the interaction between the two factors was not significant (posterior medial network: *F*(1,21) = 1.3, *p* = 0.27; regions identified using the data-driven approach: *F*(1,21) = 1.73, *p* = 0.2). These results demonstrate that objects were indeed transformed by reactivation, but the extent of object-specific transformation is comparable with the extent of the changes observed for object-context binds. This suggests that the transformation of object representations may drive the observed changes in the neural representation of object-context binds (see Discussion).

### Reactivation during sleep segregated contexts from one another

We next examined the effects of reactivation during sleep on across-context neural similarities. To test this, we correlated all objects in a cued context with all objects in other cued contexts (C↔C), as well as with all objects in non-cued contexts (C↔N). These correlations were contrasted with the correlations among objects in different non-cued contexts (N↔N; Figure 3a). If reactivation during sleep makes context more segregated from one another, we expected a decrease in correlation for the cued contexts relative to the non-cued ones. Analyses were applied both to the posterior medial network regions and to the regions identified in the data-driven approach. For the posterior medial network, a 2 x 3 repeated measures ANOVA (session x pair type) revealed no main effect of session (*F*(1,21) = 0.08, *p* = 0.78) or pair type (*F*(2,42) = 0.15, *p* = 0.86), but exposed a significant interaction between the two (*F*(2,42) = 3.6, *p* = 0.036; Figure 3b). The only contrast that emerged as significant was between C↔N and C↔C in the pre-sleep session (*t*(21) = 2.13, *p* = 0.045). When examining cross-session differences, the C↔C correlations were marginally lower than both the C↔N and N↔N correlations (*t*(21) = 2.081, *p* = 0.05; *t*(21) = 1.92, *p* = 0.07, respectively). Further analysis considered the contribution of different sub-types of object pairings (e.g., cued-to-cued between two cued contexts vs non-cued-to-non-cued between two cued contexts), revealing that effects were driven by the cued objects within the cued contexts (Supplementary Figure 3).

**Figure 3.**
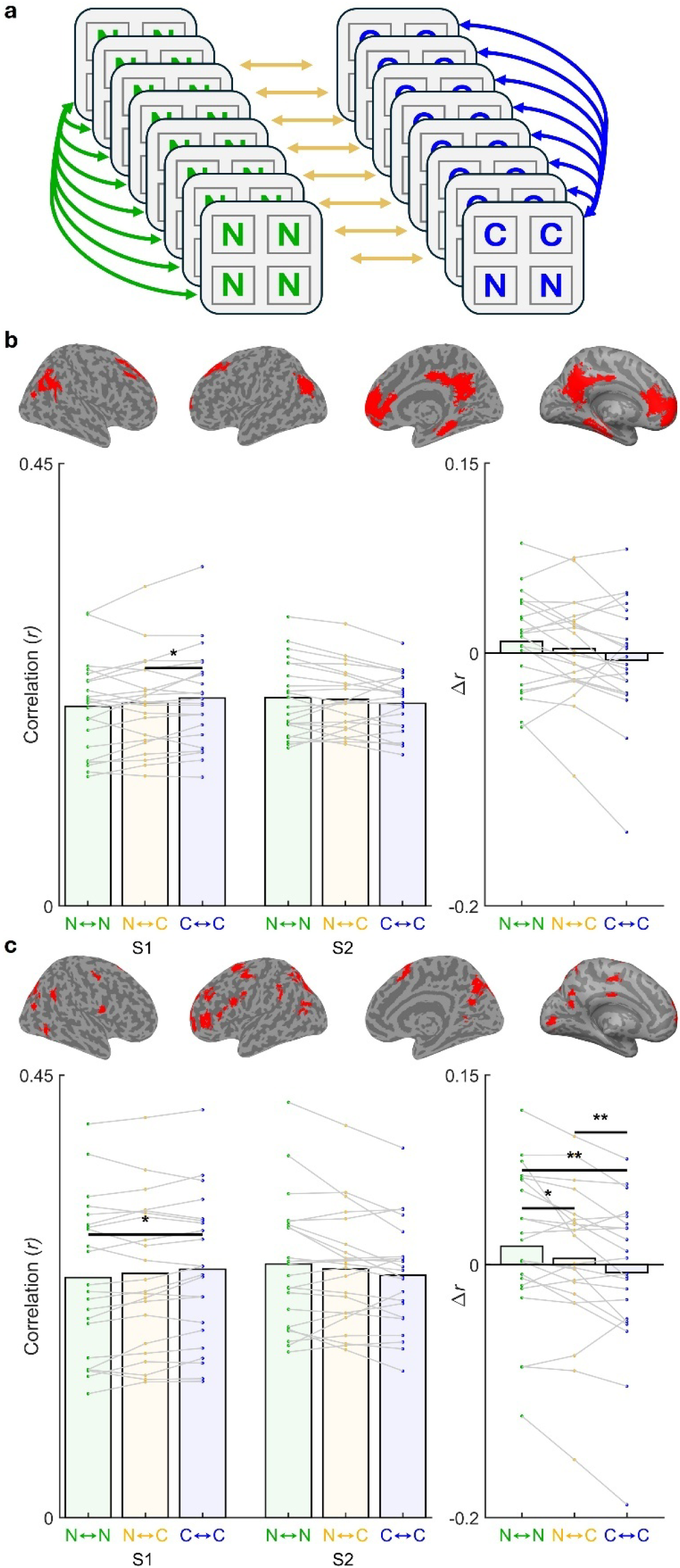
– Reactivation during sleep segregated contexts from one another. (a) Analyses compared correlations between objects across contexts, including between contexts that were both not cued

When examining the areas borne of the data-driven approach described above (Figure 2d, 2e), a clear pattern emerged across pair types (Figure 3c). The 2 x 3 repeated measures ANOVA (session x pair type) revealed no main effect of session (*F*(1,21) = 0.13, *p* = 0.72) or pair type (*F*(2,42) = 0.11, *p* = 0.89), but again, exposed a significant interaction between the two (*F*(2,42) = 8.99, *p* = 0.0006; Figure 3c). Post-hoc comparisons revealed a significant effect before sleep (N↔N < C↔C, S1, *t*(21)=2.12, *p* = 0.046) and marginal differences both before and after sleep (C↔N < C↔C, S1, *t*(21)= 1.98, *p* = 0.06; C↔C < C↔N, S2, *t*(21)= 1.86, *p* = 0.08). Critically, the cross-session changes differed for the three pair types. The correlations between different cued contexts (C↔C) decreased relative to both the cued-non-cued correlations (C↔N; *t*(21) = 2.96, *p* = 0.007) and the correlations among non-cued contexts (N↔N; *t*(21) = 3.12, p = 0.005). Furthermore, the latter two correlations differed from one another in that N↔C correlations decreased more than N↔N correlations (*t*(21) = 2.61, *p* = 0.016). As before, these effects were driven by pairing subtypes including cued objects within cued contexts (Supplementary Figure 3). Taken together, these results suggest that cuing segregated contexts from one another. Reactivation rendered the cued context more separated and less correlated. Similarly, reactivation reduced correlations between cued context and their non-cued counterparts, albeit to a lesser degree than it segregated between two cued contexts. Therefore, the results suggest that reactivation during sleep reduces the neural overlap between different contexts, thereby potentially preserving their fidelity.

The significant difference across conditions in Session 1 (i.e., before sleep) raises the concern that the effects are driven to some degree by baseline differences (e.g., whether effects can be explained by regression to the mean). Indeed, examining Session 1 data alone using repeated measures ANOVAs reveal a trend toward a significant effect for the posterior medial regions (*F*(2,42) = 2.74, *p* = 0.08) and a significant difference for the areas identified using the data-driven approach (*F*(2,42) = 3.99, *p* = 0.026). To examine whether these effects are the core driver of the interactions between session and pair type, we subsampled 17 of the 22 participants over 10,000 permutations and re-calculated the Session 1 differences and the interaction effects. Results diverged for the posterior medial regions and the data-driven regions. For both sets of regions, the permutations produced a range of *p*-values for the effect of condition in Session 1 (min = 1.17*10^-5^, max = 0.97). In the posterior medial network, most permutations led to non-significant interaction effects (36.37% of *p*s below 0.05; median *p* = 0.09), suggesting that this result is indeed less reliable and may be, at least to some extent, driven by baseline differences. When examining the permutations that resulted in no Session 1 differences using different thresholds, the interaction was not significant (e.g., when considering only permutations in which the *p*-values for Session 1 were *p* > 0.05, *p* > 0.25, *p* > 0.5, and *p* > 0.75, the average *p*-values for the session x pair type interaction were 0.16, 0.25, 0.37, and 0.52, respectively). However, in the data-driven regions, most permutations resulted in a significant interaction effect (98.51% of *p*s below 0.05; median *p* = 0.0036), suggesting that results for these regions are unlikely to be entirely due to baseline differences. When examining the permutations that resulted in no Session 1 differences using different thresholds, the interaction was consistently significant (e.g., when considering only permutations in which the *p*-values for Session 1 were *p* > 0.05, *p* > 0.25, *p* > 0.5, and *p* > 0.75, the average *p*-values for the session x pair type interaction were 0.01, 0.013, 0.013, and 0.01, respectively). In 100% of permutations, the direction of the effect was the same as reported for the full dataset (i.e., N↔N > C↔N > C↔C for Δ*r*).

All the observed differences between the pre-sleep and post-sleep sessions may have been impacted to a certain degree by cross-session differences in measurement sensitivity (i.e., changes in noise ceilings). To rule out this option, we calculated the noise ceilings by first identifying each object’s neural representation for each session and run, and then calculating the cross-run correlations between each object for each session. Results confirmed that there was no increase in the overall noise ceiling when considering the posterior medial network (*t*(21) = 0.74, *p* = 0.47) or the subregions identified in the data-driven approach (*t*(21) = 1.28, *p* = 0.21). Therefore, any differences between sessions are unlikely to account for general differences in signal-to-noise ratios across sessions.

### EEG-derived reactivation metrics predicted reductions in object-context binding

Finally, we tested whether measures of neural reactivation during sleep were related to the observed neural transformation. We first tested whether the number of cues presented during NREM correlated with the neural segregation within or between contexts. Across both the posterior medial network and the areas identified using the data-driven approach, cross-participant correlations between the number of cues and the changes in representation were not significant (*p*s > 0.55). Next, we used the EEG activity collected during sleep to examine post-sound changes in spectral activity. Previous studies have shown that post-sound changes in sigma activity (12-18 Hz), putatively reflecting sleep spindles, and delta-theta activity (2-8 Hz), putatively reflecting slow-waves and K-complexes, correlate with behavioral changes and are therefore likely to reflect neural reactivation^39–41^. Using a data-driven approach, we found two contiguous clusters in time-frequency space which correspond to those observed in previous work (*p* < 0.05, corrected; Figure 4a). Post-sound changes in power in the sigma and delta-theta ranges demonstrate that object-specific sound cues were processed by the sleeping brain.

**Figure 4.**
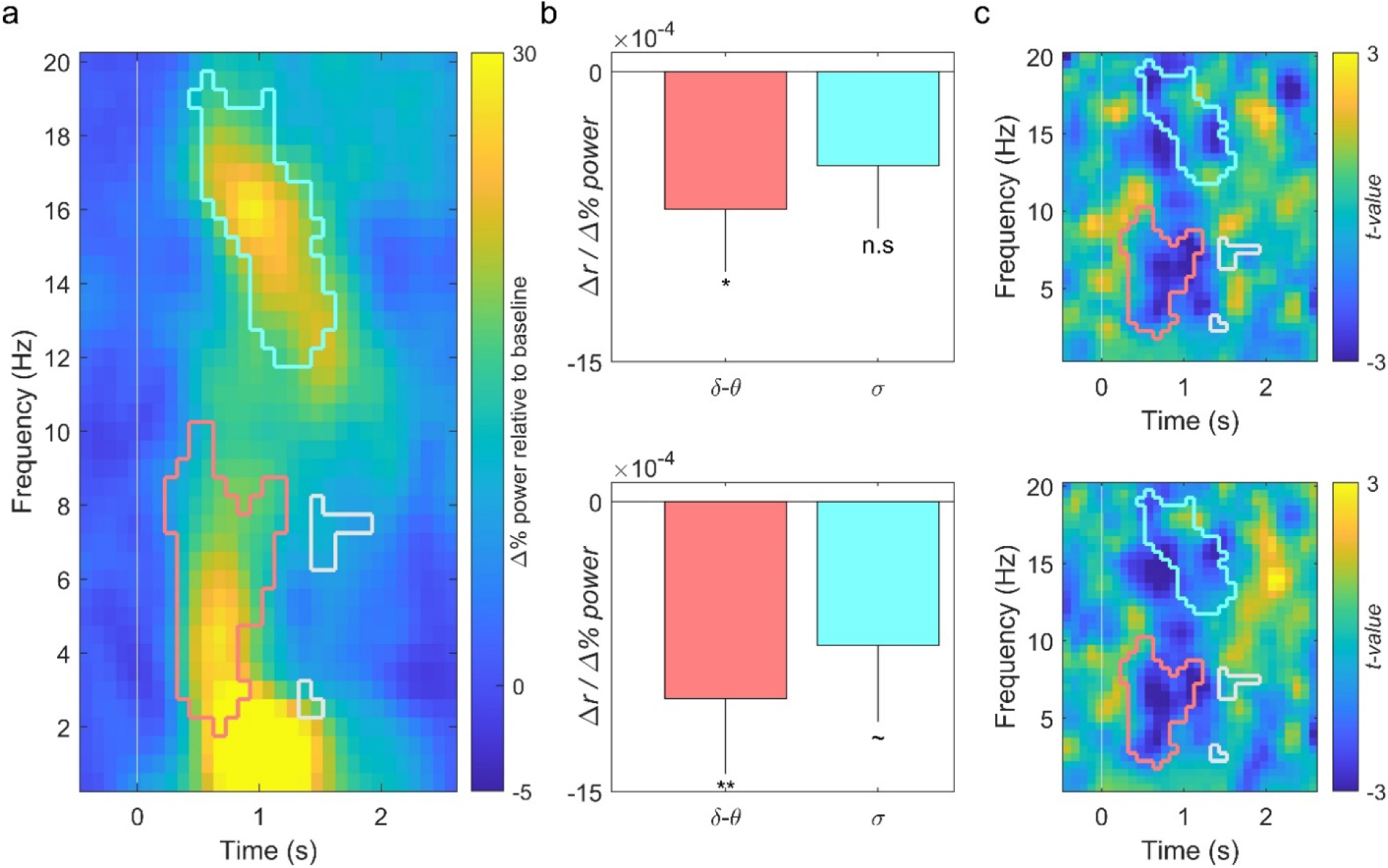
– EEG-derived reactivation metrics predicted reductions in object-context binding. (a) Sounds presented during sleep (at time 0) led to an increase in spectral power in two main clusters: one in the delta-theta range and one in the sigma range. Outlines reflect the clusters of significant points in time frequency space (p < 0.05, corrected). (b) Power changes in the clusters correlated with decreased within-context correlations among the neural correlates of objects. The top panel presents the coefficient values for the posterior medial network; the bottom panel presents the coefficient values for the areas identified in the data-driven analysis. Colors correspond to those in panel a. * p < 0.05, ** p < 0.01, ∼ p < 0.1. (c) Uncorrected t-values for correlations between the power and the within-context segregation for each point in time-frequency space. Top panel: posterior medial network; bottom panel: data-driven analysis. Colors correspond to those in panel (a).

To test whether these metrics of reactivation are related to changes in neural representations, we derived, for all sounds linked with all contexts, a measure of change in spectral power for each of the two clusters. We then tested whether changes in power predicted changes in object-context binding, which was operationalized as the correlations among the neural representations of the four contextually bound objects. This analysis was run considering the posterior medial network regions of interest (Figure 4b, top), as well as the area identified using the data-driven approach (Figure 4b, bottom). When considering the posterior medial network, there was a negative effect of delta-theta power (β = -7 x 10^-4^ Δ*r* / Δ% power, *p* = 0.028), but no significant effect of sigma power (β = -5 x 10^-4^ Δ*r* / Δ% power; *p* = 0.14). The negative coefficient indicates that higher power predicts lower correlations (i.e., more segregation), in line with the general TMR effect found in previous analyses. Similarly, for the data-driven regions, there was a significant effect of delta-theta power (β = -10 x 10^-4^ Δ*r* / Δ% power, *p* = 0.01), and a trend toward an effect of sigma power (β = -7 x 10^-4^ Δ*r* / Δ% power; *p* = 0.06). In general, activity across the frequency range around 0.5-1 s after sound onset was associated with enhanced within-context segregation across regions (Figure 4c).

## Discussion

We examined how the neural representations of memories transform when they are reactivated during sleep, focusing specifically on item-context binding. Using multiple analytical approaches and across various brain regions, we found that reactivation during sleep transforms the neural representation of objects, thereby reducing the overlap in representation between contextually bound objects. In other words, reactivation seems to segregate these memories from one another rather than enhance their shared representation. Two complementary approaches were used to identify brain regions of interest. First, we used a region-of-interest approach, focusing primarily on the posterior medial network, which has been implicated in representing contexts^35,36^ (data for other regions are shown in Supplementary Figure 1). In addition, we used a data-driven approach (corrected for multiple comparisons) to identify regions that represent contextual information in our task. Changes in delta-theta activity following sound onset during sleep, putatively reflecting memory reactivation, correlated with the within-context representational segregation.

The literature on sleep’s effects in decontextualization vs strengthening of item-context binding is mixed. Behavioral studies have demonstrated that sleep strengthens these bindings. Using multiple tasks, sleep has been shown to improve the ability to correctly identify the context in which a memory was encoded (e.g., the background image presented along with it) more so than an equally long period of wakefulness^42–45^. Furthermore, sleep deprivation has been shown to impair the binding of items to their contexts^46^. On the other hand, and as detailed above, sleep has been deemed essential for decontextualization and semantization of memories^5^. Indeed, several studies have shown that sleep is critical in generalization from previously learned exemplars to new events (e.g., ^47,48^). Generalization (i.e., creating decontextualized representations of newly learned information) and specificity (i.e., preserving contextual information linked with episodic memories) don’t necessarily compete, and both could be supported by sleep^49,50^. Unlike most previous studies, our study considered this question through the prism of neural representations rather than memory performance.

Along with the changes we observed between objects learned in the same context, we also found that reactivation impacted the degree of overlap between objects across different contexts, at least in the areas identified using the data-driven approach. Interpretation of these results was challenged by the fact that there were baseline differences (i.e., higher correlations between cued objects in different contexts in Session 1). To address this, we used a subsampling approach to show that the effect observed in the areas identified using the data-driven approach remains robust regardless of baseline differences. Our results therefore suggest that reactivation drives contexts apart from other contexts; objects in contexts that were reactivated exhibited less overlap with objects in other reactivated contexts relative to non-reactivated contexts. These results could be interpreted as an enhancement of pattern separation across contexts^51^. Behaviorally, there is evidence that sleep enhances (or at least stabilizes) pattern separation^52–54^. In terms of neural representations, the effects of sleep have not been as clear, with similar changes observed following a delay including sleep and wakefulness^52^. Critically, our study is unique in its causal approach for selectively manipulating consolidation of certain objects, suggesting that reactivation causally leads to pattern separation. These effects are observed across several cortical regions, in line with recent literature on the role of the cortex in pattern separation^55^.

Although our experimental design and analyses focus on the neural correlates of item-context binding, many other aspects of the neural underpinnings of memories are likely transformed via reactivation. Indeed, our results show that the representations of objects in cued contexts undergo more changes relative to those of objects in non-cued contexts. We demonstrate that reactivation segregated object representations from their contextual counterparts more so than from objects in other contexts. This suggests that reactivation selectively transforms item-context bindings. However, it does not imply that reactivation-induced transformations are limited to item-context binding, and it is likely that object representations transform in other manners that are beyond the scope of this manuscript. Future research should probe this process further (e.g., using representational similarity analysis, as done recently using EEG^38,56^).

Unlike previous studies employing TMR (see ^27^ for meta-analysis), our goals were to examine changes in neural representations rather than changes in retrieval likelihood. Classical models of consolidation, such as the Complementary Learning Systems model^5^, focus primarily on changes in neural representation (i.e., change from hippocampal dependency to cortical dependency), yet this type of transformation has rarely been directly examined. Our design, therefore, intentionally overtrained participants, resulting in retrieval performance at ceiling levels, and therefore no effect of TMR on memory performance. This is not a limitation of our design, but rather a feature of it; if changes over the delay included not only changes in an object’s neural representation but also changes in the probability of retrieval, those two effects would be confounded. For example, if a memory was unlikely to be retrieved before sleep, and the retrieval probability increased after sleep, the BOLD signal could reflect this change in retrieval probability rather than differences in the functional neuroanatomy of item-context binding, and these two effects would be hard to disentangle and examine separately. The literature on consolidation predicts changes in neural representations over sleep and time, and this study was aimed at measuring those rather than changes in memory performance.

Having said that, a limitation that arises from this design choice is that there is no behavioral evidence for the benefits of TMR in this study and using this paradigm. Performance remained at ceiling levels even one week after learning. However, the EEG data collected while participants slept provide complementary evidence for the effectiveness of the TMR manipulation: the observed activity patterns mirror those presented in previous TMR studies (e.g., ^39–41^), and the correlations between EEG reactivation patterns and fMRI-derived changes in representation converge with our main findings to demonstrate that our manipulation was impactful. Another limitation of our design is that the neural representations of contexts were not examined directly, but rather operationalized as the representational overlap among contextually bound items. Indeed, other studies have used a common visual scene to monitor context reinstatement (e.g., ^19,57,58^). Our definition of context was intended to be broader (e.g., an imagined episode) rather than linked to a specific visual representation, but this design choice limited our ability to directly monitor context reinstatement. Finally, the study lacks a control group that either remained awake or slept without cues being presented. This limits the interpretation of our results, as it is not entirely clear whether the changes observed for the non-cued objects were due to the passing of time, the natural effects of sleep, or some unintended effects of the cuing procedure applied to the other objects.

Of note, the effects of reactivation during sleep on memories within their contexts have recently been studied by examining changes in memory performance, demonstrating that the behavioral effects of reactivation are similarly guided by contexts^59,60^ (see also ^61^). Curiously, however, these results demonstrated that memory benefits spill over to contextually related memories during sleep. At face value, these results seem to be contradictory, showing that reactivation benefits contextually related memories on the one hand, while weakening item-context binding on the other (see also ^50^). However, these results are not necessarily in conflict. For example, it may be that contextual reinstatement during sleep itself leads to decontextualization. Future studies should continue to probe the relationship between these effects, as well as the relationship between neural and behavioral manifestations of decontextualization.

The main results concerning the effects of reactivation on item-context binding were observed both in the posterior medial network and in a set of regions that represented the narrative contexts used in this study, identified using a data-driven approach. Our a-priori hypothesis was that context would be represented by the nodes of the posterior medial network, as shown in previous work (e.g., ^35,36,62^). Indeed, these regions did reflect context information in our task (i.e., showed higher within-context vs between-context correlation patterns). However, surprisingly, there is little overlap between the nodes of the posterior medial and the areas identified using the data-driven approach. The source of this discrepancy is unclear and may have to do with unique aspects of our experimental design or with the task demands in the scanner. Specifically, our operationalization of context diverges from many other studies, which themselves differ from one another (see discussion in ^63^). Our task included participant-generated narratives in which objects shared an imagined time, location, and causal relationship. However, these narratives were not experienced directly, unlike contexts employed in most other studies (e.g., video clips depicting episodes^35^, objects presented alongside a shared scene^19^). Nevertheless, it is unclear why this design would impact the functional neuroanatomy supporting contextual information.

Taken together, our results provide evidence that reactivation during sleep transforms the neural representations of objects, rendering then more distinguishable from their contextual counterparts. As a result, reactivation decreases neural overlap both within contexts and across contexts, potentially leading to both decontextualization and pattern separation (Figure 5). These findings constitute an important step toward understanding sleep’s role in the reorganization of memory representations. Future work should consider longer periods of time, rather than a single bout of sleep, as has been done in other studies that did not focus exclusively on sleep (e.g., ^17,18^). Despite the clear appeal of using causal manipulations of sleep, recently developed approaches for examining endogenous reactivation events during sleep provide the analytical foundation to further exploration of how sleep transforms memory reactivation^64,65^. Using a combination of causal and observational techniques would lead to a better understanding of sleep’s role in shaping neural representations, not only for objects in context, but for all forms of memory as they evolve over time.

**Figure 5.**
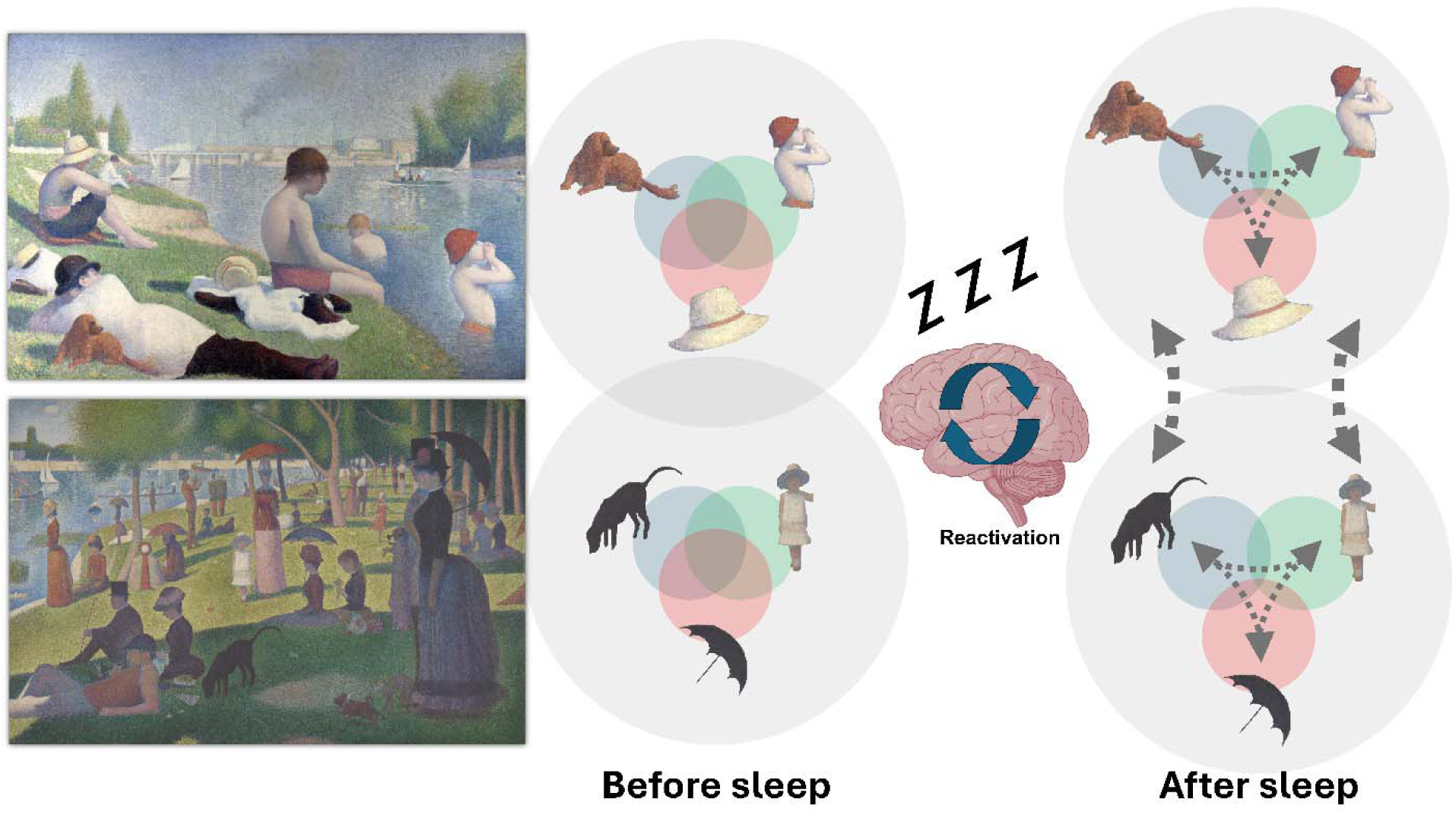
– Conceptual recap of main results. When experiencing an event, episodic memories include multiple elements that are bound together by a context (e.g., bathing in the river, top left; a picnic at the park, bottom left). The neural representations supporting memory for these elements overlap, reflecting their shared context (center). During sleep, some of these memory traces are reactivated, leading to two main consequences: a segregation between contextually bound elements (triangular thin arrows); and a segregation between the contexts themselves (vertical thick arrows). This figure uses work by Georges Seurat (Bathers at Asnières, top; A Sunday Afternoon on the Island of La Grande Jatte, bottom), which are in the public domain.

## Materials and Methods

### Participants

In total, 26 participants were recruited for this study (22.8 ± 0.64 SEM years old, 19 women and seven men). Potential participants completed a screener to confirm that they are eligible for MRI studies. Those who had metal implants, were claustrophobic, or were pregnant were excluded from participation. Participants were asked to go to bed later in the night before the study, wake up earlier on the morning of the study, and avoid any caffeine on the day of the study. Four participants were excluded from the final dataset: Two participants were not sufficiently exposed to sounds during sleep (one was not exposed to any sounds, the other was not exposed to sounds during non-REM sleep), one participant quit mid-study due to claustrophobia, and one participant was excluded due to poor performance in the task (> 8 standard deviations worse than other participants). The remaining 22 participants were 22.59 ± 0.71 years old and included 17 women and five men. All participants consented to participate in the experiment. The protocol was approved by the Institutional Review Board at Northwestern University.

### Materials

Stimuli consisted of 68 square images of objects (e.g., animals, household items), taken primarily from the Bank of Standardized Stimuli^33^. Some images were taken from online image databases (e.g., freeimages.com). Each image was paired with a congruent sound (e.g., cat - “meow”). Sounds were mostly taken from online sources (e.g., freesound.org). On average, these sounds were 520 ms long (SD = 93 ms; max length = 605 ms). An additional set of 64 images was used as foils in the follow-up task. These images were also taken from the Bank of Standardized Stimuli or online sources.

Stimuli were presented using different devices in different phases of the experiment. During training and testing outside of the MRI scanner, sounds and images were presented using a laptop computer (Lenovo TR460s, Hong Kong, China). During MRI scanning sessions, images were presented on a rear projection screen (NordicNeuroLabsLCD 3.0.5), and sounds were presented through in-ear headphones (Sensimetrics model S14, transmitting the sound with Avotec SiletScan model ss-3000). During sleep, sounds were presented using speakers placed near the participants’ heads (Rev A00, Harman Kardon, CT, USA). All in-lab tasks were coded in MATLAB (version 2020b, MathWorks Inc., MA, USA), using the Psychophysics Toolbox^66^. The online portion of the task was coded using Qualtrics (Qualtrics Inc., WA, USA).

### Procedure

The experiment included two parts, set 7-9 days apart (Figure 1a). The first part lasted approximately six hours and was completed in the lab. After completing a consent form and a safety questionnaire, participants began a training session, followed by an MRI session. Then, participants were fitted with an EEG cap and napped for approximately 90 minutes outside of the scanner. Next, they returned for another scanning session, followed by a final testing session outside of the scanner. The second part, which lasted approximately 15 minutes, was completed online 7-9 days after the first. Participants used their own computers for this part.

### Training session

During the training session, participants were first presented with scenarios (e.g., “Getting a haircut”, “Attending a birthday party”) along with four images of objects (Figure 1b). Object-scenario associations were randomized across participants. The task was to craft a unique and memorable narrative incorporating the objects in the provided setting. Participants were guided to tell the story in the first person, make it creative, imaginative, and bizarre, and keep it contained within the provided scenario (i.e., avoid changing context mid-story). In total, participants formed 17 stories, including an initial practice story. Participants were exposed to each scenario title along with images and labels for the four objects. Before forming the story, they were presented with the congruent sounds linked with each object by clicking on its image. Then, after they had formed each story in their minds, they recorded an audio clip in which they described it, before moving on to the next scenario. All steps were self-paced.

After recording all stories, participants practiced the Retrieval Task (Figure 1c). This was the main task used in both MRI scanning sessions, intended to evoke associative memories between scenarios and objects. In each trial, participants were presented with one of the images for 4 s, accompanied by the object’s congruent sound. The instructions were to bring to mind which scenario the object was linked with. On some trials, termed No-Response Trials, memory for the association was not tested; the image was followed by an intertrial interval, in which a black cross was presented on screen (mean duration = 2 s, range 1-4 s). On other trials, termed Response Trials, memory for the association was tested; after image offset, four optional scenarios were presented on screen (e.g., “Haircut”, “Birthday”). Participants had 3.75 s to indicate the correct scenario by pressing a button using one of four fingers placed on the keyboard. No feedback was given, and after the 3.75 s time window ended, the intertrial interval began. The rationale for using both trial types was practical. Using only Response Trials, which would have provided a stronger behavioral index, would have extended each in-scanner block (see below) from 8.5 minutes to ∼11 minutes, increasing fatigue and reducing data quality. To minimize scanning time and participant demands, we therefore included Non-Response Trials. Critically, when an image was presented, and throughout the first 4 s of the trial, participants had no way to know whether a trial would be a Response or No-Response Trial and were asked to think of the correct scenario as fast as possible to prepare for a rapid response. At the end of each run of the task, the percentage of correct responses was presented to the participant.

In the training session, a short practice version of the Retrieval Task was used. This version of the task included eight trials and consisted of the four objects associated with the practice scenario, each presented twice in a random order. Half of the trials were Non-Response Trials, and half were Response Trials. Responses were made on four designated keyboard keys. Participants were informed that this task would be repeated in the scanner and that they are expected to retrieve all formed associations, not just the practice ones. After completing the Retrieval Task, participants were given an opportunity to improve their memory for the stories they constructed. The purpose of this task was to bring associative memories close to ceiling levels. Participants were sequentially exposed to each scenario in a random order and asked to list the associated objects. If they had trouble remembering all four, they were allowed to listen to a recording of their own story until they clearly remembered all objects that were associated with it. For each scenario, participants recorded their responses (i.e., a spoken list of the four objects) using a microphone. All steps were self-paced.

The final training task was on a Perceptual Task, which was also used in the scanner. Originally, this task was intended to detect non-task-related activation of contexts upon exposure to the objects. However, data for this task were not incorporated in any analyses included in this manuscript and will likely be analyzed separately ahead of a future publication. The task is still described here for completeness. For this task, participants were exposed to images of all 68 objects sequentially. Some images were partially occluded by a small gray rectangle, which covered ∼2% of the image. Participants had to indicate, using one of two buttons, whether the image was occluded by the rectangle or not. Each image was presented for 2 s, followed by an intertrial interval in which a black cross was presented (mean duration = 2 s, range 1-4 s). At the end of the task, the percentage of correct responses was presented. The practice version of this task, used for training, included two presentations of each of the four practice objects, presented in random order. Half of the trials included images occluded by the rectangle. After completing this task, participants prepared for the MRI session.

### First MRI session

Scanning was done at Northwestern University’s Center for Translational Imaging, using a Siemens 3T Prisma whole-body scanner. Participants lay in the scanner, were fitted with in-ear headphones covered by noise-blocking earmuffs and handed a four-button controller. The volume was adjusted to the participants’ comfort, while ensuring that sounds were audible over scanner noise. Then, participants underwent an anatomical T1-weighted scan (3D Multi-Echo (ME) MPRAGE sequence TR = 2300 ms, TE = 2.38 ms, inversion time (TI) = 902 ms, slices per slab = 240, resolution = 0.8 x 0.8 x 0.8 mm^3^, flip angle = 8°). During the scan, they were completed the two practice tasks mentioned above again (the practice versions of the Retrieval Task and Perceptual Task).

After the anatomical scan, participants began five runs of the memory-association task, during which functional MRI scans were collected. (1700 ms TR, 32 ms TE, flip angle = 75°, 82 slices, resolution = 2.0 x 2.0 x 2.0 mm^3^). Each block consisted of 66 trials, starting with two trials presenting practice objects. One of these first trials was a Response Trial, and the other was a Non-Response Trial. The remaining 64 trials included all the objects incorporated in the 16 stories. Forty percent of these trials were Response Trials. Each image was presented once in each of the five runs, so that, for each image, two runs included Response Trials and three runs included Non-Response Trials. Each run lasted approximately 8.5 minutes, and participants were offered short breaks in between runs. Participants’ performance was continuously monitored to confirm that they remained awake and engaged and that they avoided excessive movement.

The last task completed in the scanner was the Perceptual Task. A single run of this task was presented, consisting of 68 trials, with the first four trials including practice objects. Half of the objects were obscured by a rectangle. This run lasted approximately 4.5 minutes. Participants then left the scanner.

### Afternoon nap

Participants next prepared for a 90-minute EEG-monitored afternoon nap (Figure 1d). During sleep, 16 different object-related sounds were repeatedly presented unobtrusively. The cued objects were strategically chosen so that memory for half of the objects (two objects) in half of the stories (eight stories) would be reactivated during sleep (Figure 1e).

Participants were first fitted with a 32-channel EEG cap, along with two reference electrodes placed on the mastoids, two electrooculogram electrodes placed around the eyes, and one electromyography electrode placed on the chin (BrainVision Inc. NC, USA). Next, participants slept on a bed in a darkened room. White noise was presented through speakers placed near the participant’s head. EEG data were monitored online by an experimenter. When participants entered slow-wave sleep, the experimenter began the cuing protocol. Sounds were presented continuously, at a stimulus onset asynchrony jittered between 4.5 and 5.5 s. Each 16-sound sequence included all sounds in a randomized order. Sound presentation was discontinued upon arousal or transition into rapid-eye-movement sleep. If participants did not reach slow-wave sleep 45 minutes into the nap, sounds were presented either during slow-wave sleep or stage 2 of sleep during the remainder of the 90-minute nap. Two participants who were not exposed to each of the 16 sounds at least once during these stages of sleep were excluded from analysis. Table 2 shows the distribution of cues over sleep stages, as assessed offline. Participants were awoken after approximately 90 minutes and were allowed to wash their hair.

**Table 2.** Sleep architecture and cue-presentation statistics across stages (average ± SEM)

|  | Wake | N1 | N2 | N3 | REM |
| --- | --- | --- | --- | --- | --- |
| **Minutes** | 18.80 ± 3.06 | 17.23 ± 1.81 | 33.45 ± 2.68 | 16.93 ± 3.22 | 3.43 ± 1.4 |
| **% of time in bed** | 20.65% ± 1.84% | 18.98% ± 1.84% | 37.15% ± 2.82% | 18.5% ± 3.53% | 4.72% ± 2.23% |
| **Average # of sounds** | 2.86 ± 2.49 | 2.68 ± 1.41 | 59 ± 10.73 | 99.23 ± 22.2 | 0.18 ± 0.18 |

### Second MRI session

The second MRI session began shortly after participants were awoken (mean delay = 37 minutes, range 23-55 minutes). The second session was similar to the first. This time, however, volume was preset to the level established beforehand rather than adjusted based on participant preference. Additionally, the session did not include an anatomical scan. Participants were offered the opportunity to engage in the practice versions of the Retrieval and Perceptual Tasks (*n* = 3 completed the first, *n* = 2 completed the second). Then, they began five full runs of the Retrieval Task, followed by one run of the Perceptual Task. In both tasks, objects were presented in a different pseudorandom order, following the guidelines described for the first MRI session.

### Final testing session

After leaving the scanner, participants engaged in one final testing session. In this session, they were presented with each of the 16 scenarios and asked to list the four related objects using the microphone (Recall Task). This task was self-paced. After this task, participants were debriefed. They were asked whether they remembered hearing any sounds during sleep. Three participants who mentioned hearing task-related sounds then underwent an additional task, in which they were asked to indicate, for each sound, whether they remembered it being played during sleep. Performance in this task was close to chance level (12.5%, 25%, 31.25% of presented sounds were correctly identified; chance = 25%). Participants were given instructions for the follow-up tests and dismissed.

### Follow-up tests

Participants were tested on their memories online 7-9 days after completing the first part of the experiment. This part consisted of two tasks. The first task, a Recognition Task, was implemented on Qualtrics, and required participants to determine whether a set of images was old (learned in the previous sessions) or new. A total of 132 objects were presented in random order, including four practice images, 64 previously learned objects, and 64 novel objects. After completing this task, participants were asked to sort the objects based on the scenario with which they were matched. For this Sorting Task, a Google Slides document was shared with the participants, which included images of all 68 objects arranged on the left part of the slide (in a 4 x 17 array occupying ∼20% of the slide). The slide also included 17 boxes, each labeled with the name of a scenario, along with a text label with instructions (“Please drag each item on the left to the box with the title of the story”). Participants had to drag the images of all 68 objects to their correct boxes (e.g., if the Ball was linked with the Birthday scenario, they would need to drag the Ball image to the box labeled Birthday for it to be scored correct). We did not record the order of object placement, but rather the accuracy of the end result. Both tasks were self-paced. Upon completion of these tasks, participants were compensated.

### Behavioral analyses

Behavioral data were analyzed using MATLAB 2020b (MathWorks Inc., Massachusetts). Data from the Retrieval Task were tabulated from Response Trials for each object and session, across runs, and were then averaged across objects and submitted into repeated measures ANOVA. As performance for all subsequent memory tests (the Recall Task, Recognition Task, and Sorting Task) was at ceiling levels, only descriptive statistics were calculated. For the Recall and Sorting Tasks, the number of correctly identified/sorted objects was tabulated, whereas for the Recognition Task, both hits and false alarms were calculated and reported. Practice contexts were not considered in these analyses.

### EEG preprocessing and sleep scoring

EEG and electrooculography data were filtered between 0.3 and 35 Hz and electromyography data were filtered between 10 and 100 Hz. Electrodes with consistently noisy data were interpolated. The resulting dataset was then sleep-scored by two trained researchers in 30-s epochs and consolidated by a third researcher. Scorers were not exposed to information regarding the timing of sound presentations.

### MRI data preprocessing

Data include one T1-weighted (T1w) image and 10 BOLD runs per subject (across all tasks and sessions). Preprocessing was performed using fMRIPrep 20.2.1^67^ (RRID: SCR_016216), which is based on Nipype 1.5.1^68^ (RRID: SCR_002502). More information is provided in the Supplemental Materials.

### fMRI Data Analysis

General linear models (GLMs) were run in AFNI using the 3dDeconvolve function. Regressors for framewise displacement, cerebrospinal fluid, white matter, and head motion (six directions plus their six derivatives) were included as predictors of no interest. Two different sets of GLMs were run, each employing the Least Squares – Separate approach^69^ and producing a β-map that was used in subsequent analyses. The first set of GLMs, used for most analyses reported in the manuscript, was run for each object across all runs within a session (64 GLMs total per session; GLMs were not run for practice objects). Each GLM included one predictor for the specific object (across its five presentations, locked to stimulus onset), one predictor for all other objects (67 objects across five runs), and one predictor for button responses. The second set of GLMs was used for the noise ceiling analysis. GLMs were run for each object and each run (64 x 5 = 320 GLMs total per session). Each GLM included one predictor for the specific object in a specific run (a single event, locked to stimulus onset), one predictor for all other object-presentation events (67 objects across five runs and the target object across four runs), and one predictor for button responses. Both Response and Non-Response trial data were used for both analyses. The β maps produced by the cross-run GLMs were used to examine similarities between different objects, based on their contextual associations (Figure 2a, Figure 3a). These analyses were done using MATLAB 2020b (MathWorks Inc., Massachusetts). Only objects that were correctly associated with their context in the Retrieval Task (in both Session 1 trials) were used for analysis. In our analyses, the cross-run β maps for each object were correlated with the other objects within its own context or with objects in other contexts using Pearson’s *r*. In the correlation analyses run across sessions, we correlated the cross-run β maps for each object in session 1 with the cross-run β maps in session 2. These analyses either correlated an object’s own representation across sessions, or the representation of an object in one session with the representation of its contextual counterparts in the other session. All correlations were z-transformed, and z-values were averaged across objects within each participant and submitted to further statistical analyses. Averaged z-values were transformed back into *r***-**values for visualization in figures. Repeated measures ANOVAs were used to evaluate differences between conditions and sessions, followed by t-tests for pairwise comparisons.

### Regions of interest

To understand how individual memory representations were supported by core memory structures, we focused our analysis on the posterior medial network, including its subregions, which has been shown to be linked with episodic memories, and specifically contextual memories^35,36,62,70^. We defined these regions based on a cortical atlas (areas labeled under Default A and Default B in ^37^, as done in previous studies (e.g., ^71^). In addition, we examined other memory-related regions implicated in consolidation, such as the hippocampus and ventromedial prefrontal cortex (e.g., ^19^). Both regions were defined based on the Harvard-Oxford atlas^72^, and were examined separately in the left and right hemispheres to assess potential hemispheric differences in memory representation or reactivation effects. Results for these regions are included in Supplementary Figure 1. For each region of interest, we examined the extent to which it represented context in our task by correlating the β values for each object with those of other objects within the same context and statistically comparing the obtained correlation values with correlations with objects linked with other contexts.

### Data-Driven Analyses

To examine context representation across the brain without constraining the analysis to predefined regions of interest, we conducted data-driven, whole-brain analyses using two approaches: (1) a parcel-based approach, using the Schaefer 1000-parcel functional atlas analysis^37^; and (2) a searchlight approach using spheres of varying radii (3, 4, 5, and 6 voxels; Supplementary Figure 2). For the parcel-based analysis, we began by focusing on data from Session 1, prior to any cuing or sleep, to determine which parcels showed sensitivity to item-context binding. For each parcel, we correlated β values for each object with those of other objects within the same context and statistically compared the obtained correlation values with correlations with objects linked with other contexts (Figure 2d). All correlations were Pearson’s *r* and were z-transformed before averaging and comparisons. The 1-tailed t-tests yielded 1,000 *p*-values, which were corrected using the False Discovery Rate method^73^. For each searchlight analysis, a sphere was systematically moved across the brain to compute item–context correlations within each local neighborhood, as it was done for the parcels. This analysis was conducted to confirm that no subcortical regions remained unidentified using the parcellation approach, which covered only the cerebral cortex. The identified regions (*p* < 0.001, uncorrected), along with the main results for these regions, are presented in Supplementary Figure 2.

### EEG Analyses

EEG data were parsed into segments starting 2.5 s before and ending 3 s after the onset of each sound presented during slow-wave sleep or stage 2 of sleep. Segments including artifacts were excluded from further analysis. For each segment, a spectrogram was calculated using data from two frontocentral electrodes (i.e., electrodes Cz and Fz; *spectrogram* function in MATLAB; 0.75 s window, 0.65 s overlap, frequencies between 0.5 and 20 Hz in 0.5 Hz increments). Data for all segments were averaged within participant, and the frequency-specific baseline between -1.5 and -0.5 s was used for baseline correction for all trials. For each point in time-frequency space, we used a *t*-test to compare the baseline-corrected power to zero across participants (α = 0.05, Bonferroni corrected for multiple comparisons). Two contiguous clusters emerged, reflecting activity in the delta-theta and sigma ranges, akin to those identified in previous studies^39,59,74,75^. These clusters putatively reflect sound-locked increases in slow-wave activity and spindles, respectively, and have been used as a marker of neural reactivation. Power was extracted for both clusters for each baseline-reduced segment within each participant.

We hypothesized that reactivation, operationalized by power within these clusters, would lead to greater segregation within contexts (i.e., reduced correlation among neural representations of contextually bound objects). To test this, we first estimated the spectral power change linked with each context (in each cluster and for each participant). For each context, we identified all EEG segments including one of the two sounds related to it and averaged the power for these segments in each of the two clusters. This was repeated across participants, segments, and clusters, and these values acted as our independent variables. As a dependent variable, we derived a measure of within-context correlations by averaging, within each context, the pairwise correlations in neural representations among all pairs of objects. This was repeated for each context, participant, and session, using both the regions of interest described above and the regions identified using the data-driven approach. Finally, we used a mixed linear model with a random effect of participant to test whether the power predicted the change in correlation between Session 1 and 2. We repeated the same analysis for each point in time-frequency space (i.e., agnostic of the identified clusters) to prepare Figure 4c.

## Acknowledgments

ES is supported by the US National Institutes of Health [grant numbers K99-MH122663 and R00-MH122663]. The authors wish to thank Nicholas J. Lew for assistance with sleep scoring, Rachael Anne Young, Joel Voss, and Christina Zelano for assistance with data collection and equipment, and Allison Tran, Asieh Zadbood, Megan Peters, and Craig Stark for assistance with analyses. This work was supported by the NIDA IRP of the National Institute of Health (NIH) (ZIA DA000642 to TK). The contributions of the NIH authors are considered Works of the United States Government. The findings and conclusions presented in this paper are those of the authors and do not necessarily reflect the views of the NIH or the U.S. Department of Health and Human Services.

## Declaration of interests

The authors declare no competing interests.

## Supplemental Materials

**Supplementary Figure 1.**
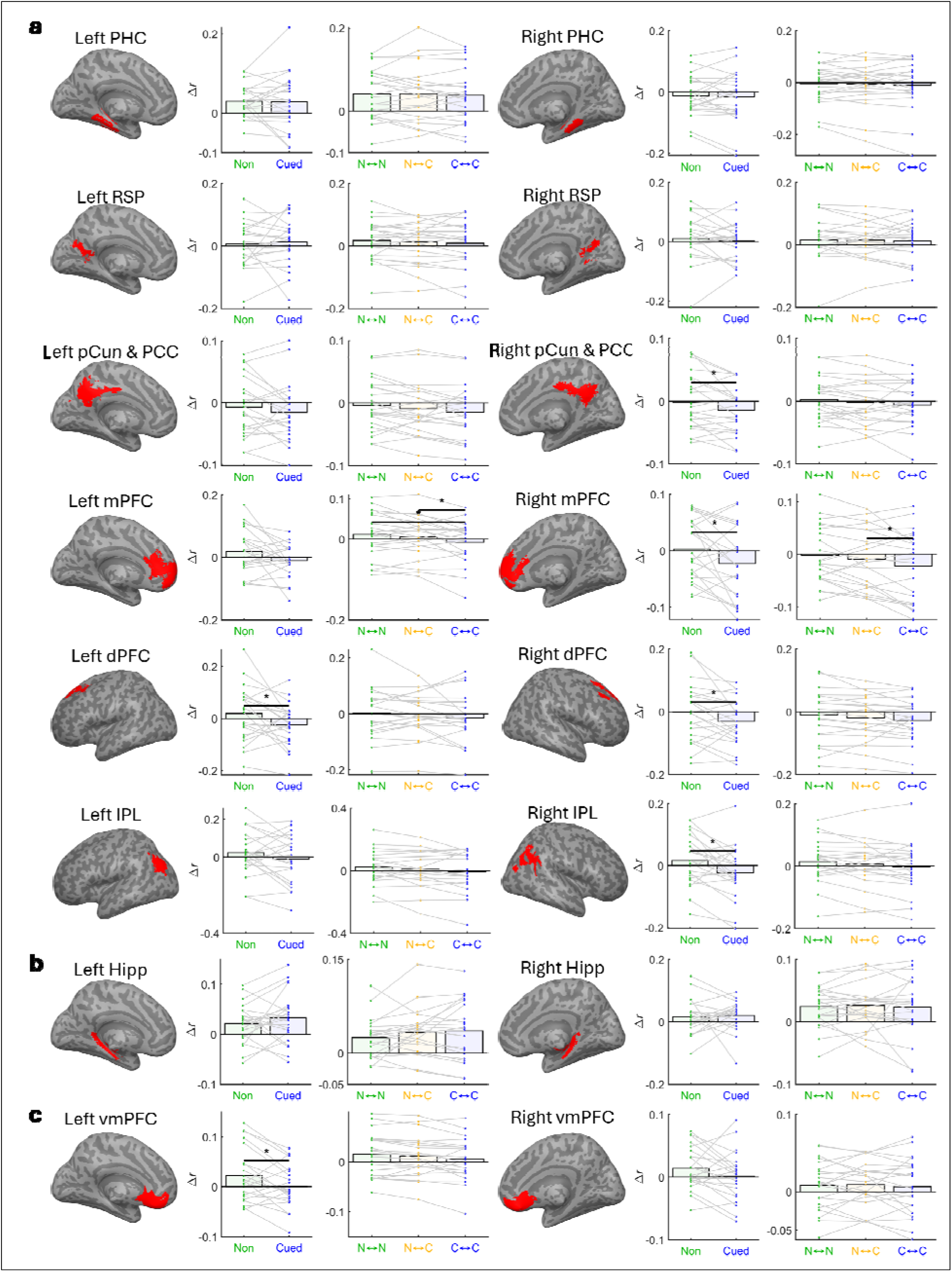
Results for regions of the posterior medial network, the hippocampus, and the ventromedial prefrontal cortex. (a) Results for different regions of the posterior medial network, shown for the left and right hemispheres. For each region, cross-session changes in correlation coefficients are shown within-context (left) and between-context (right) analyses, mirroring Figures 2 and 3, respectively. PHC – parahippocampal cortex; RSP – retrosplenial cortex; pCun – precuneus; PCC – posterior cingulate cortex; mPFC – medial prefrontal cortex; dPFC – dorsal prefrontal cortex; IPL – inferior parietal lobule. (b) Same for the hippocampus. Analyses were also done for anterior and posterior hippocampus in both hemispheres (i.e., the most anterior/posterior thirds of the coronal slices) with qualitatively similar null results (not shown). (c) Same for the ventromedial prefrontal cortex. N↔N – non-cued to non-cued contexts; C↔N – cued to non-cued contexts; C↔C – cued to cued contexts. * - p < 0.05.v

**Supplementary Figure 2.**
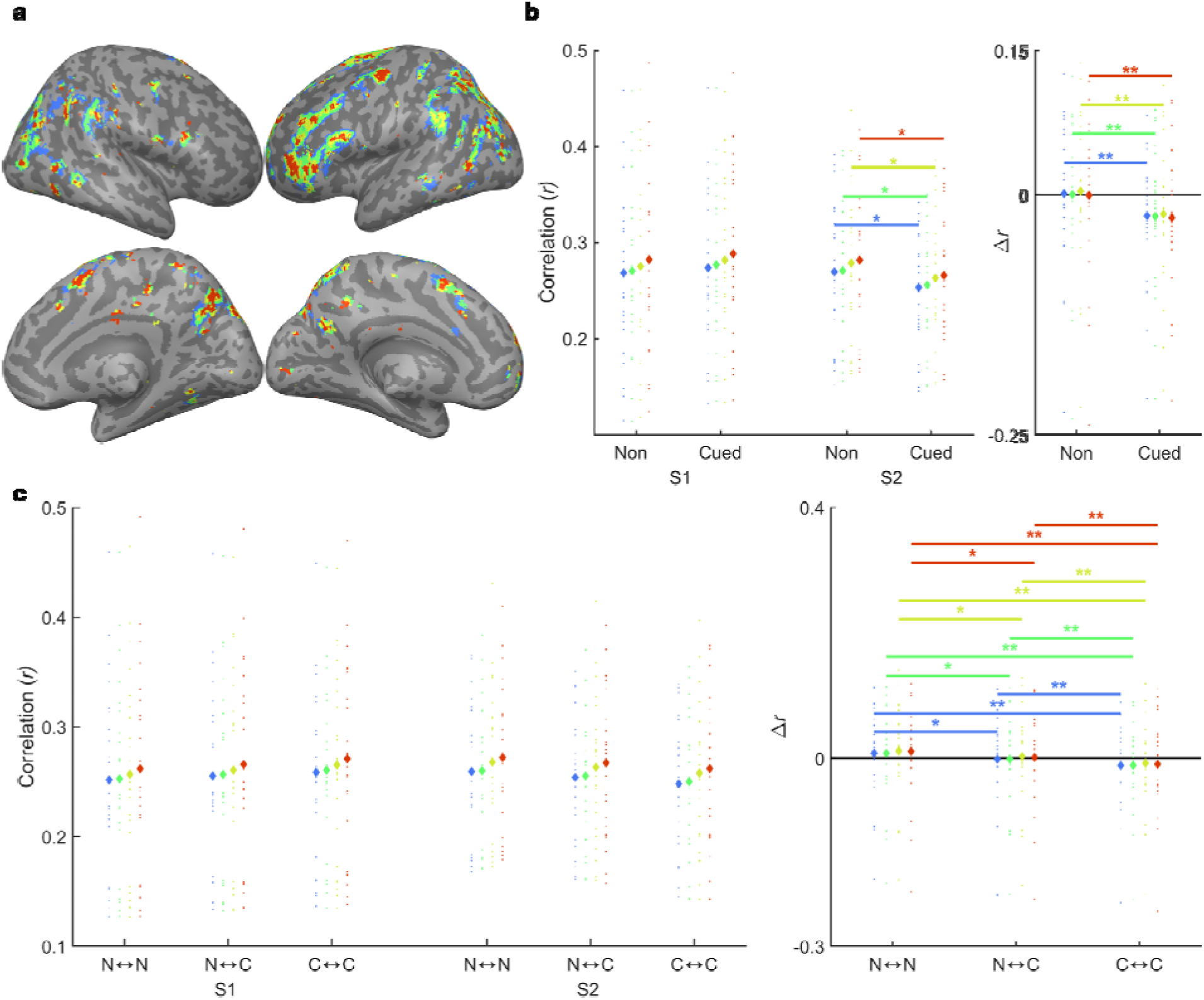
Results for context-sensitive brain regions, as revealed using searchlight analysis employing a sphere with a radius of 3, 4, 5, or 6 voxels (red, yellow, green, blue, respectively). (a) Regions identified using the searchlight approach (p < 0.001, uncorrected). Note that regions are restricted to the cortex and are qualitatively similar to those revealed using the parcel-based approach (Figure 2e, left). (b) Results for the within-context analysis (compare with Figure 2). (c) Results for the between-context analysis (compare with Figure 3). S1 – Session 1 (pre-sleep); S2 – Session 2 (postsleep); N↔N – non-cued to non-cued contexts; C↔N – cued to non-cued contexts; C↔C – cued to cued contexts. * - p < 0.05, ** - p < 0.01, *** - p < 0.001.

**Supplementary Figure 3.**
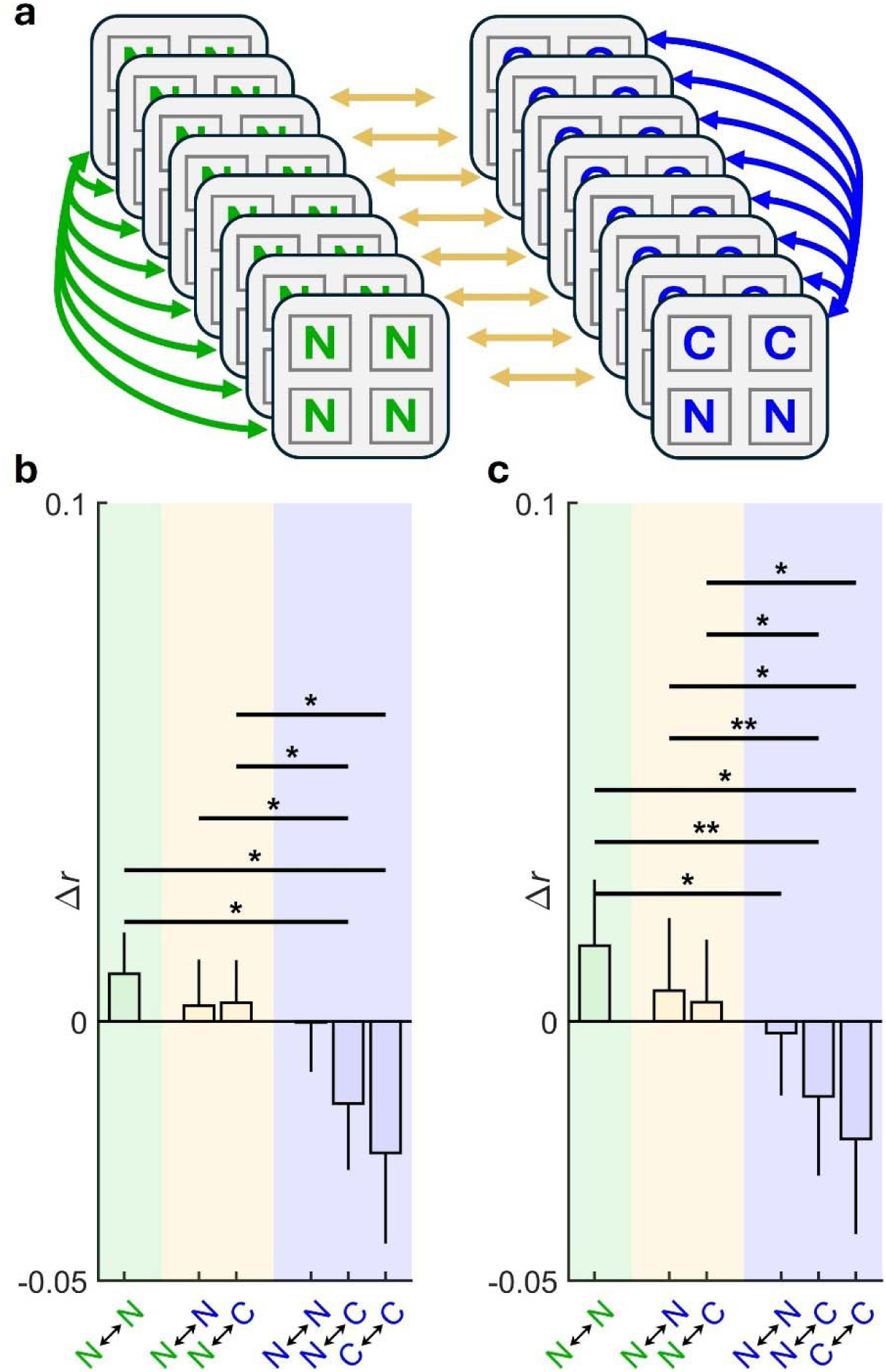
Correlations across different contexts shown for different sets of object pairings. (a) As in Figure 3, analyses compared correlations between objects across contexts, this time considering different types of pairings, as elaborated below. (b) and (c) show changes in correlation between pre- and post-sleep for the posterior medial network regions and the parcel identified using the data-driven approach, respectively. In both panels, the x-axes labels refer to (from left to right): non-cued objects in non-cued contexts ↔ non-cued objects in other non-cued contexts; non-cued objects in non-cued contexts ↔ non-cued objects in cued contexts; non-cued objects in non-cued contexts ↔ cued objects in cued contexts; non-cued objects in cued contexts ↔ non-cued objects in other cued contexts; non-cued objects in cued contexts ↔ cued objects in other cued contexts; cued *objects in cued* contexts ↔ *cued objects in other cued contexts. Error bars reflect between subject standard errors, but analyses were done within participants using paired t-tests. N – non-cued; C –cued. * - p < 0.05, ** - p < 0.01.*

**Supplementary Figure 4.**
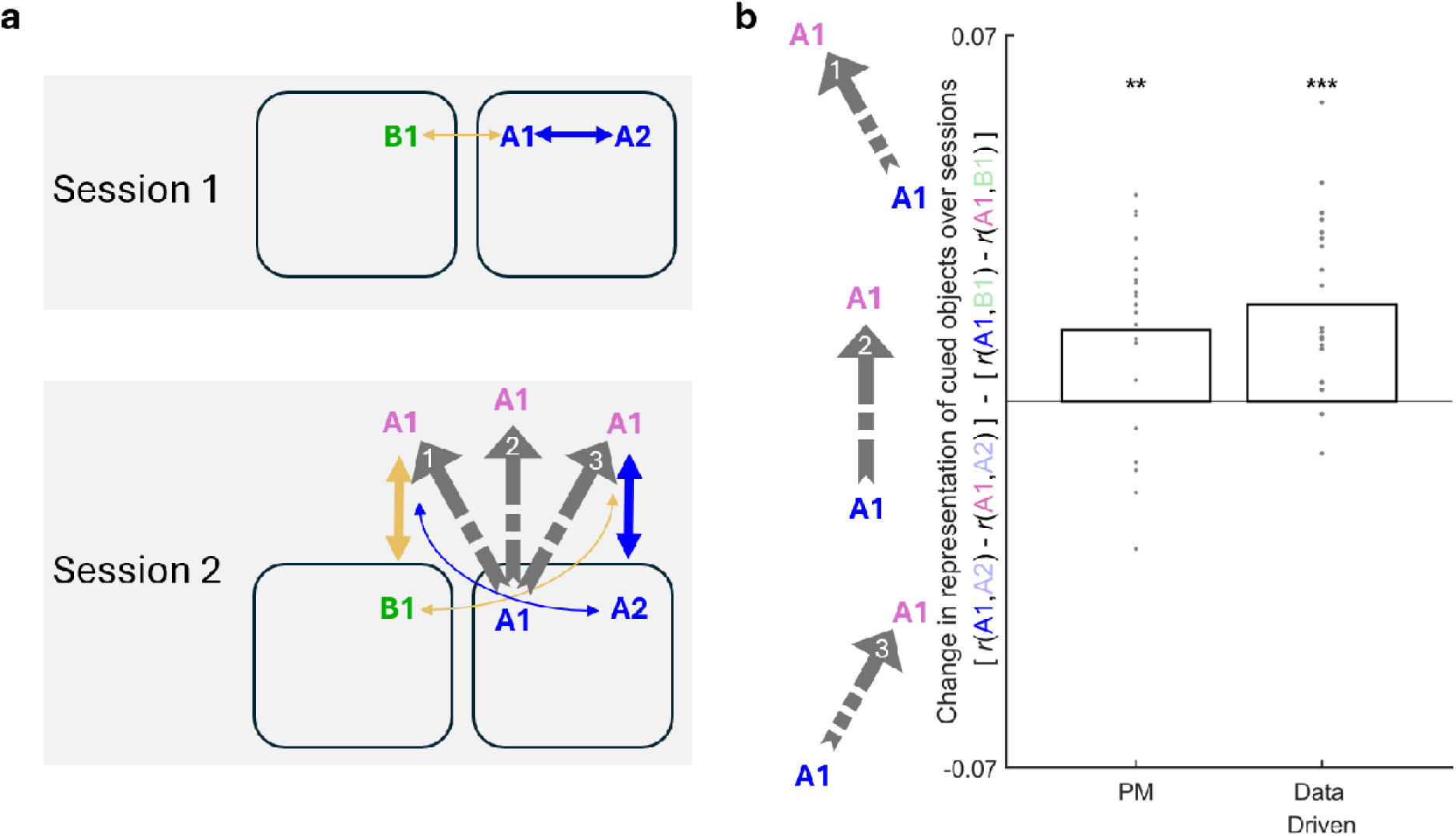
Changes in item-to-item representational correlations following reactivation are larger within- relative to between-contexts. (a) Schematic of analytic approach. We examined the correlations among objects in cued contexts (marked as A1, A2) and non-cued contexts (marked as B1). The top subpanel shows the correlations among the representations of these objects in Session 1 (thicker arrows indicate higher correlations). After reactivation, object A1 changes its representation, as reflected by the change in its location and color. The three gray arrows illustrate three hypothetical trajectories in a simplified representational space. Arrow 1 shows a transition that renders A1’s new representation closer to B1 and further from A2 (i.e., reactivation led to higher within- relative to between-context correlations). Arrow 3 shows the opposite trend, and Arrow 2 reflects a scenario whereby A1’s change in representation was independent of context altogether. (b) Results for the PM networks and for areas identified using the data-driven approach presented in Figure 2d. Colors in the y-label follow panel (a). The gray arrows illustrate that higher values reflect a shift toward higher within- relative to between-context correlations. PM – posterior medial network. ** - p < 0.01, *** - p < 0.001.

## Supplemental Methods

### Anatomical data preprocessing

A total of 1 T1-weighted (T1w) images were found within the input BIDS dataset. The T1-weighted (T1w) image was corrected for intensity non-uniformity (INU) with N4BiasFieldCorrection (Tustison et al. 2010), distributed with ANTs 2.3.3 (Avants et al. 2008, RRID:SCR_004757), and used as T1w-reference throughout the workflow. The T1w-reference was then skull-stripped with a Nipype implementation of the antsBrainExtraction.sh workflow (from ANTs), using OASIS30ANTs as target template. Brain tissue segmentation of cerebrospinal fluid (CSF), white-matter (WM) and gray-matter (GM) was performed on the brain-extracted T1w using fast (FSL 5.0.9, RRID:SCR_002823, Zhang, Brady, and Smith 2001). Brain surfaces were reconstructed using recon-all (FreeSurfer 6.0.1, RRID:SCR_001847, Dale, Fischl, and Sereno 1999), and the brain mask estimated previously was refined with a custom variation of the method to reconcile ANTs-derived and FreeSurfer-derived segmentations of the cortical gray-matter of Mindboggle (RRID:SCR_002438, Klein et al. 2017). Volume-based spatial normalization to two standard spaces (MNI152NLin2009cAsym, MNI152NLin6Asym) was performed through nonlinear registration with antsRegistration (ANTs 2.3.3), using brain-extracted versions of both T1w reference and the T1w template. The following templates were selected for spatial normalization: ICBM 152 Nonlinear Asymmetrical template version 2009c [Fonov et al. (2009), RRID:SCR_008796; TemplateFlow ID: MNI152NLin2009cAsym], FSL’s MNI ICBM 152 non-linear 6th Generation Asymmetric Average Brain Stereotaxic Registration Model [Evans et al. (2012), RRID:SCR_002823; TemplateFlow ID: MNI152NLin6Asym]

### Functional data preprocessing

For each of the 10 BOLD runs found per subject (across all tasks and sessions), the following preprocessing was performed. First, a reference volume and its skull-stripped version were generated using a custom methodology of fMRIPrep. A deformation field to correct for susceptibility distortions was estimated based on fMRIPrep’s fieldmap-less approach. The deformation field is that resulting from co-registering the BOLD reference to the same-subject T1w-reference with its intensity inverted (Wang et al. 2017; Huntenburg 2014). Registration is performed with antsRegistration (ANTs 2.3.3), and the process regularized by constraining deformation to be nonzero only along the phase-encoding direction, and modulated with an average fieldmap template (Treiber et al. 2016). Based on the estimated susceptibility distortion, a corrected EPI (echo-planar imaging) reference was calculated for a more accurate co-registration with the anatomical reference. The BOLD reference was then co-registered to the T1w reference using bbregister (FreeSurfer) which implements boundary-based registration (Greve and Fischl 2009). Co-registration was configured with six degrees of freedom. Head-motion parameters with respect to the BOLD reference (transformation matrices, and six corresponding rotation and translation parameters) are estimated before any spatiotemporal filtering using mcflirt (FSL 5.0.9, Jenkinson et al. 2002). The BOLD time-series were resampled onto the following surfaces (FreeSurfer reconstruction nomenclature): fsnative. The BOLD time-series (including slice-timing correction when applied) were resampled onto their original, native space by applying a single, composite transform to correct for head-motion and susceptibility distortions. These resampled BOLD time-series will be referred to as preprocessed BOLD in original space, or just preprocessed BOLD. The BOLD time-series were resampled into several standard spaces, correspondingly generating the following spatially-normalized, preprocessed BOLD runs: MNI152NLin2009cAsym, MNI152NLin6Asym. First, a reference volume and its skull-stripped version were generated using a custom methodology of fMRIPrep. Several confounding time-series were calculated based on the preprocessed BOLD: framewise displacement (FD), DVARS and three region-wise global signals. FD was computed using two formulations following Power (absolute sum of relative motions, Power et al. (2014)) and Jenkinson (relative root mean square displacement between affines, Jenkinson et al. (2002)). FD and DVARS are calculated for each functional run, both using their implementations in Nipype (following the definitions by Power et al. 2014). The three global signals are extracted within the CSF, the WM, and the whole-brain masks. Additionally, a set of physiological regressors were extracted to allow for component-based noise correction (CompCor, Behzadi et al. 2007). Principal components are estimated after high-pass filtering the preprocessed BOLD time-series (using a discrete cosine filter with 128s cut-off) for the two CompCor variants: temporal (tCompCor) and anatomical (aCompCor). tCompCor components are then calculated from the top 2% variable voxels within the brain mask. For aCompCor, three probabilistic masks (CSF, WM and combined CSF+WM) are generated in anatomical space. The implementation differs from that of Behzadi et al. in that instead of eroding the masks by 2 pixels on BOLD space, the aCompCor masks are subtracted a mask of pixels that likely contain a volume fraction of GM. This mask is obtained by dilating a GM mask extracted from the FreeSurfer’s aseg segmentation, and it ensures components are not extracted from voxels containing a minimal fraction of GM. Finally, these masks are resampled into BOLD space and binarized by thresholding at 0.99 (as in the original implementation). Components are also calculated separately within the WM and CSF masks. For each CompCor decomposition, the k components with the largest singular values are retained, such that the retained components’ time series are sufficient to explain 50 percent of variance across the nuisance mask (CSF, WM, combined, or temporal). The remaining components are dropped from consideration. The head-motion estimates calculated in the correction step were also placed within the corresponding confounds file. The confound time series derived from head motion estimates and global signals were expanded with the inclusion of temporal derivatives and quadratic terms for each (Satterthwaite et al. 2013). Frames that exceeded a threshold of 0.5 mm FD or 1.5 standardised DVARS were annotated as motion outliers. All resamplings can be performed with a single interpolation step by composing all the pertinent transformations (i.e. head-motion transform matrices, susceptibility distortion correction when available, and co-registrations to anatomical and output spaces). Gridded (volumetric) resamplings were performed using antsApplyTransforms (ANTs), configured with Lanczos interpolation to minimize the smoothing effects of other kernels (Lanczos 1964). Non-gridded (surface) resamplings were performed using mri_vol2surf (FreeSurfer).

Many internal operations of fMRIPrep use Nilearn 0.6.2 (Abraham et al. 2014, RRID:SCR_001362), mostly within the functional processing workflow. For more details of the pipeline, see the section corresponding to workflows in fMRIPrep’s documentation.

## Notes

### Competing Interest Statement

The authors have declared no competing interest.

### Summary of Updates

New results motivated by review comments, including anaysis of EEG-based reactivation measures.

